# DIMERBOW: exploring possible GPCR dimer interfaces

**DOI:** 10.1101/836213

**Authors:** Adrián García-Recio, Gemma Navarro, Rafael Franco, Mireia Olivella, Ramon Guixà-González, Arnau Cordomí

## Abstract

G protein-coupled receptors (GPCRs) can form homo-, heterodimers and larger order oligomers that exert different functions than monomers. The pharmacological potential of such complexes is hampered by the limited information available on the type of complex formed and its quaternary structure. Several GPCR structures in the Protein Data Bank display crystallographic interfaces potentially compatible with physiological interactions. Here we present DIMERBOW, a database and web application aimed to visually browse the complete repertoire of potential GPCR dimers present in solved structures. The tool is suited to help finding the best possible structural template to model GPCR homomers. DIMERBOW is available at http://lmc.uab.es/dimerbow/.

## Introduction

5-10% of genes in the human genome code for environment-sensing cell surface receptors, most of which are G protein-coupled receptors (GPCRs). This family of receptors respond to a huge variety of stimuli, including ions, amino acids, lipids, peptides, proteins and even photons. For long, GPCRs were described as monomeric transmembrane (TM) receptors, but it is now well accepted that they can form homo- and heteromer complexes in cells (Gurevich and Gurevich, 2018; Borroto-Escuela et al., 2014; Fuxe et al., 2010). They constitute novel signaling units with unique functional and regulatory properties that are opening new opportunities for drug discovery (Ferré, 2015; Fuxe et al., 2010; Franco et al., 2016). While GPCR heteromers have been thoroughly characterized by biophysical and biochemical methods (Sleno and Hébert, 2018; Xue et al., 2019; Guo et al., 2017), the structural basis behind the allosteric communications between receptor protomers remains poorly understood. Most GPCRs interact via their TM domain, except for class C GPCRs, where both extracellular and TM domains are involved (Koehl et al., 2019). Several crystal structures of GPCRs reveal crystallographic dimers compatible with the spatial constraints imposed by the membrane (Cordomí et al., 2015; Katritch et al., 2013; Simpson et al., 2010). Despite some of these dimers might not represent biologically relevant interfaces outside the crystal lattice, they are certainly valuable templates to model GPCR homomers and heteromers. Stenkamp identified 71 parallel dimeric crystallographic interfaces (Stenkamp, 2018) in a systematic analysis of the Protein Data Bank (PDB) (Berman et al., 2000). Additionally, molecular dynamics (MD) simulations of crystallographic dimers have also been used to shed light on the structural basis of GPCR homodimers (Baltoumas et al., 2016; Johnston and Filizola, 2014). Here we present DIMERBOW, a web application (available at http://lmc.uab.cat/dimerbow) aimed at exploring the large and growing repertoire of crystallographic dimers of GPCRs involving the TM domain. DIMERBOW features a systematic analysis of protomer-protomer contacts along with the results from coarse-grained MD simulations of all feasible dimers. The database currently contains information from 97 interfaces and represent a unique tool to model GPCR dimers and oligomers. An automated pipeline for processing new interfaces, submitting simulations and performing data analysis guarantees an up-to-date resource.

## Methods

### Database of putative dimers

DIMERBOW relies on a database of putative GPCR dimers (Suppl. Note 1) from protomer-protomer pairs observed in structures deposited in the PDB (Berman et al., 2000).

### Coarse-grained MD simulations

To study the feasibility of crystallographic interfaces, we used coarse-grained MD simulations with the MARTINI v2.1 force field (Marrink et al., 2007). We simulated a total of 97 dimeric interfaces totaling to 300 μs (Suppl. Note 2). All simulations were run using GROMACS v.2018.5 (Abraham et al., 2015).

### Implementation and update

DIMERBOW relies on a Python (v.3.7) backend that uses the Flask framework (v.1.0.2). Dimer data is stored in a MySQL database (v.8.0.18). Interactive plots and molecule visualizations employ Bokeh (v.1.3.5b) and NGL (v.2.0.0, (Rose and Hildebrand, 2015)), respectively. Preparation of experimental dimeric structures, submission of MD simulation and analysis pipelines are all automated using Python, Bash and R scripts. This guarantees regular updates as new structures become available.

## Results

DIMERBOW offers two main functionalities: i) browse putative dimers from high-resolution structures of GPCRs deposited in the PDB (Fig. 1), and ii) examine the results of MD simulations of the putative dimers inserted in a model POPC membrane (Suppl. Fig. S1). In both browsers, a central panel features one (reference) protomer surrounded by the second protomers displayed as circles. Mouse actions on these circles can open an interactive 3D representation of each dimer in the right panel or display extra information including the number of residue-residue contacts, helices involved, and dimer stability in MD simulations. Finally, a toolkit in the left panel allows filtering by receptor type, helices involved, minimal number of interactions in the dimer interface, or interaction mode.

**Fig. 1.**
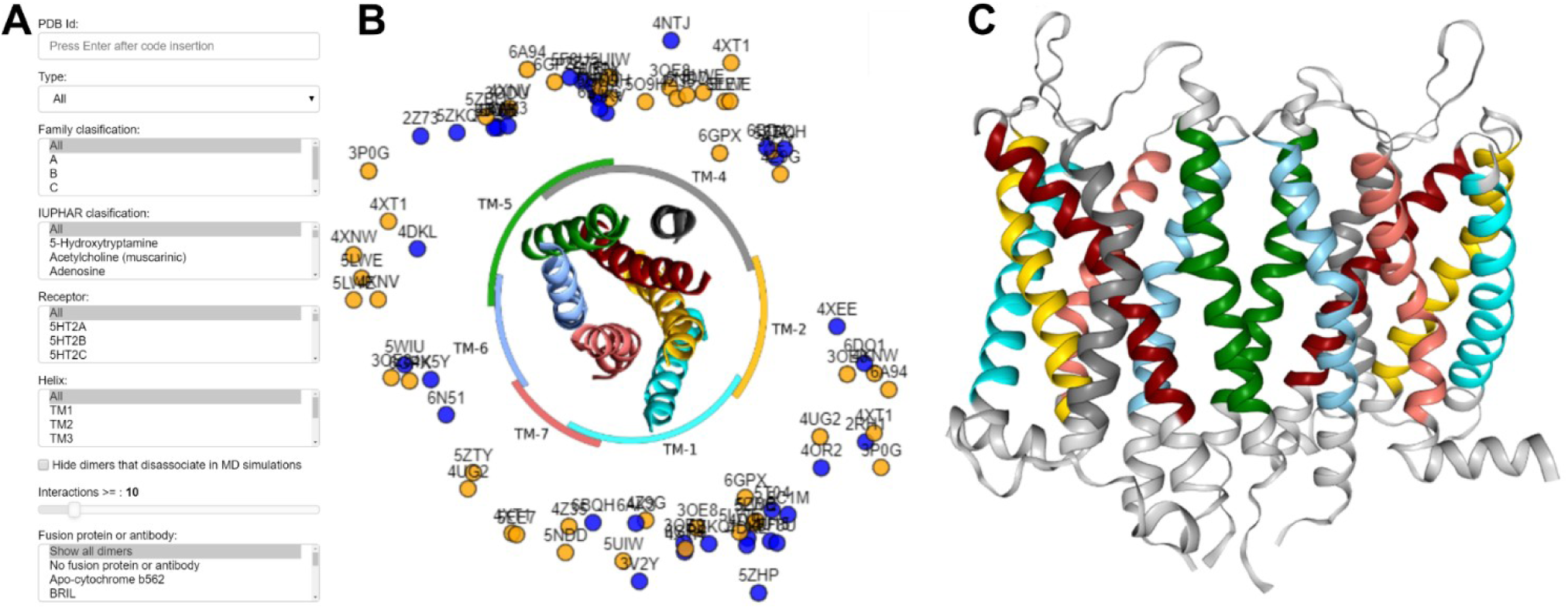
The PDB dimer browser. **A**) filtering options, **B**) central panel displaying a schematic representation of all dimers, and **C**) interactive visualization of each dimer structure in NGL (Rose and Hildebrand, 2015).

Suppl. Table 1 summarizes the number of dimeric interfaces present in the current release. We used MD simulations (see Methods) to assess the stability of these dimers outside the crystal lattice and embedded in a model lipid bilayer. 53 out of 97 dimers (55%) did not dissociate during the simulations and, overall, crystallographic dimers with larger number of interactions are more stable (Suppl. Note 3). Regarding dimer’s symmetry a larger number of head-to-tail (HT) dimers dissociated when compared to head-to-head (HH) dimers (Suppl. Table 1). In line with previous reports (Guo et al., 2008; Liang et al., 2003; Katritch et al., 2013), a significant number of dimers involve TM1 (29%), TM4 (19%), and TM5 (26%) (see Suppl. Table 1). HH dimers tend to involve TM1 together with TM2 or TM7, and TM5 together with TM4 or TM6, whereas HT dimers mostly involve TM1 interacting with TM4.

## Conclusion

DIMERBOW is a web application for rapid, intuitive, and systematic visualization of the complete repertoire of crystallographic interfaces for GPCR homodimers available in the PDB. The tool is suited to help computational scientists, molecular biologists and crystallographers interested in GPCR dimers and oligomers finding the best possible structural template to model GPCR homomers or heteromers or to ascertain how frequent a specific dimeric interface occurs. The tool has the potential to be expanded to other families of membrane proteins forming dimers or oligomers.

## Supporting information

Supplementary inormation

