## Supplementary inormation for "DIMERBOW: exploring possible GPCR dimer interfaces"

### Supplementary Methods

#### Supplementary Note 1: Database preparation

In its current version (30/03/2019) we processed 213 G protein-coupled receptors (GPCR) structures that contained the PFAM (Finn et al., 2016) domains PF00001, PF00002, PF00003 and PF01534 that correspond to the transmembrane (TM) signature of class A, B, C or F GPCRs, respectively. For each Protein Data Bank (PDB) structure, we generated all crystallographic protomer-protomer contacts (pairs) using the generate *symmetry mates* function of PyMol v2.1.1 (https://pymol.org). We only kept pairs where both protomers were close (centers-of-mass <48Å), parallel (angle between the principal axis of both protomers <30º) and compatible with the membrane plane (i.e. difference between the Z-direction components of their centers-of-mass <10Å according to the Orientations of Proteins in Membranes database (Lomize et al., 2012)). For each dimer, fusion proteins and nanobodies (often added to help the crystallization) were removed together with any non-protein molecule (water, lipids, …) at >8Å from any protomer. Since the purpose of DIMERBOW is to explore TM interactions, we removed the large and flexible N-terminal domains present in class B, C and F. For proteins where different structures have been solved (same Uniprot accession code) with a similar interface (root-mean-square-distance (rmsd) <3Å after superposition of Cα), we used the structure with the best resolution. The classification in subfamilies is taken from (Harding et al., 2018)

#### Supplementary Note 2. Set-up an MD simulation protocol

Each crystallographic dimer was embedded in a lipid bilayer comprised of ~600 1-palmitoyl-2-oleyl-phosphatidyl-choline (POPC) molecules that was subsequently solvated with water and 0.15 M NaCl. The martinize.py (de Jong D.H. et al., 2013) and insane.py (Wassenaar, T.A. et al.,2015) scripts were used to generate the coarse-grained topology and structure files. To keep the overall shape of individual protomers we employed an ElNeDyn elastic network (Periole et al., 2009). Each system was subjected to energy minimization, followed by multi-step equilibration (16.5 ns in total) and production (600 ns) runs. We simulated five replicas of each system.

#### Supplementary Note 3. Stability of dimers in MD simulations

While in our simulated dimers tend to dissociate or decrease in number of contacts between protomers (see main text and Supplementary Table 1), in some simulations we observe an increase in the number of interactions. Different aspect of the new environment created around the protein during the set-up of the simulations (see Supplementary Note 2) may be behind this effect, including i) the release of restrictions imposed by crystal packing, ii) the removal of crystallographic fatty acids, detergents and lipids upon system preparation, enabling direct contacts between promoters, and/or iii) the removal of crystallographic stabilizer proteins.

**Supplementary Table 1. Summary count of dimers in DIMERBOW current release**
PDB dimers are listed by (A) number of residue-residue interactions (without any filtering), (B) preservation of the dimeric structure in MD simulations (**B**), TM helix or helices involved (**C**). The number of dimers is counted separately for head-to-head and head-to-tail modes (for protomers with and without two-fold rotational symmetry; see Stenkamp, 2018). Cumulative numbers are shown in the rightmost column. We chose a conservative criterion for tagging a receptor as preserved in MD simulations consisting in average RMSD ≤ 1.40Å for the best 3 out of 5 replicates. Despite it is generally assumed that GPCRs dimerize through head-to-head (HH) interfaces, the current release of DIMERBOW displays a non-negligible 40% of dimers in the head-to-tail (HT) mode.

| **(A)** | **Head-to-head** | **Head-to-tail** | **Total** |
|---|---|---|---|
| All dimers | 59 | 38 | **97** |
| ≥10 interactions | 38 | 23 | 61 |
| <10 interactions | 21 | 15 | 36 |

| **(B)** | **Head-to-head** | **Head-to-tail** | **Total** |
|---|---|---|---|
| Preserved in MD | 34 | 19 | 53 |
| Dissociate in MD | 25 | 19 | 44 |

| **(C)** | **Head-to-head** | **Head-to-tail** | **Total** |
|---|---|---|---|
| TM1 | 17 | 11 | 28 |
| TM2 | 9 | 4 | 13 |
| TM3 | 6 | 4 | 10 |
| TM4 | 8 | 10 | 18 |
| TM5 | 14 | 11 | 25 |
| TM6 | 3 | 5 | 8 |
| TM7 | 4 | 6 | 10 |

**Supplementary Figure S1. The MD simulations browser.**

For each dimer, the MD browser displays **A**) a panel with selection and filtering options, **B**) a central panel with a schematic representation of the last snapshots of each trajectory (at 600 ns) compared to the starting dimer from the solved structure, and **C**) a panel with an interactive visualization of the last snapshot of any trajectory using NGL (Rose and Hildebrand, 2015). In **B**, the center of mass of each MD replicate is shown in gray whereas the initial PDB structure is shown in green. When MD brings asymmetry in the dimer structure each simulation is shown twice (one dot per possible superposition using the first or the second protomer).

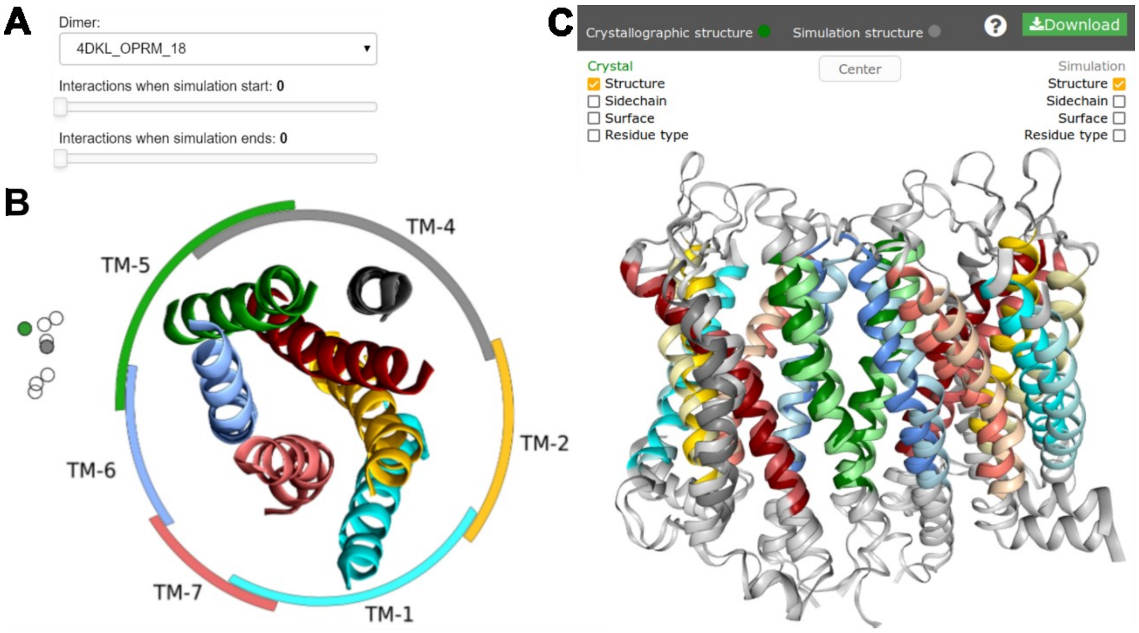
